# Structural characterization of a Thorarchaeota profilin indicates eukaryotic-like features but with an extended N-terminus

**DOI:** 10.1101/2021.08.06.455371

**Authors:** Raviteja Inturi, Sandra Lara, Mahmoud Derweesh, Celestine N. Chi

## Abstract

The emergence of the first eukaryotic cell was preceded by evolutionary events which are still highly debatable. Recently, comprehensive metagenomics analysis has uncovered that the Asgard super-phylum is the closest yet known archaea host of eukaryotes. However, it remains to be established if a large number of eukaryotic signature proteins predicated to be encoded by the Asgard super-phylum are functional at least, in the context of a eukaryotic cell. Here, we determined the three-dimensional structure of profilin from Thorarchaeota by nuclear magnetic resonance spectroscopy and show that this profilin has a rigid core with a flexible N-terminus which was previously implicated in polyproline binding. In addition, we also show that thorProfilin co-localizes with eukaryotic actin in cultured HeLa cells. This finding reaffirm the notion that Asgardean encoded proteins possess eukaryotic-like characteristics and strengthen likely existence of a complex cytoskeleton already in a last eukaryotic common ancestor.

## INTRODUCTION

The emergence of the first eukaryotic cell was preceded by evolutionary events, which are still highly debatable^*1-4*^. However, it is now generally accepted that these events led to the merger between an archaeal host and an alphaprotobacterium. The Asgards (Lokiarchaeota, Thorarchaeota, Odinarchaeota, Heimdallarchaeota, Wukongarchaeota, Hodarchaeota, Kariarchaeota, Hermodarchaeota, Gerdarchaeota and Baldrarchaeota), a newly discovered superphyla within the archaea has emerged as the last eukaryotic common ancestor (LECA)^*5-8*^. The Asgard genomes encode numerous eukaryotic signature proteins previously unseen in prokaryotes ^*5,7*^. These proteins are important for regulation of cell processes such as cell motility, information transfer, ribosomal, trafficking and ubiquitin system. In eukaryotes motility is highly driven by actin, a key cytoskeletal protein whose dynamics is regulated by many other adapter proteins such as gesolin, profilin, vasodilator-stimulated phosphoprotein (VASP), actin related protein (ARP) 2/3 complex and a handful of signaling molecules including (Phosphatidylinosi-tol-4,5-bisphosphate (PIP_2_))^*9-12*^. The structures of profilins from several members of the Asgard super-phyla including Lokiarchaeota, Odinarchaeota and Heimdallarchaeota have been determined at high resolution both by NMR and X-ray crystallography^*13,14*^. These structures revealed that Asgard encode a typical eukaryotic-like profilin fold. Furthermore, these profilins were shown to regulate eukaryotic actin polymerization *in vitro* and this process was modulated by profilin interaction with PIP_2_^*13,14*^. In addition, it was found that some members of the Asgard archaea (including Heimdallarchaeota LC3 phyla) encode profilins with an atypical N-terminus extension that modulates their interaction with polyproline, an interaction not observed in members of the Loki and Odin phyla^*13*^. Interestingly, sequence and phylogenetic analysis data revealed that certain members including Thorarchaeota encode profilins with an extended N-terminal region^*14*^. However, sequence alignment alone does not prove if the Thorarchaeota profilins do indeed have the profilin fold and if they have any functional role. Here, we show that Thorarchaeota (*A0A524EIQ6_THOAR*) a candidate phylum within the Asgard superphylum, encodes a putative profilin (thorProfilin) with a eukaryotic fold and co-localizes with F-actin filaments *in vivo* indicating a complex evolutionary relationship within the Asgardarchaea and reaffirming the central role in eukaryogenesis within the Asgardian.

## RESULTS

### Thorarchaeota archaeon profilin has a typical profilin but with an N-terminal extension

The recent comprehensive metagenomics analysis that uncovered the Asgard super-phylum as the closest yet known archaea host of eukaryotes was the major breakthrough in the study of eukaryogenesis^1,2^. However, it remains to be established if a large number of eukaryotic signature proteins (ESPs) predicated to be encoded by the Asgard super-phylum are functional^1,2^. Asgards encode profilins with a very low sequence conservation and as such the true nature of the profilins can only be ascertain via structural homology. Structures of several profilins including Loki-1, Loki-2, Odin and Heimdall LC3 have been determined previously by X-Ray crystallography both individually and bound to rabbit actin^*13*^ or by NMR^*14*^. As a first step towards establishing the identity of the Thorarchaeota profilin, we determined the three-dimensional structure of Thorarchaeota profilin with uniprot accession number *TFG12995*.*1* by nuclear magnetic resonance spectroscopy. First, we performed a complete NMR assignments of the thorProfilin using a series of uniform and selective amino acid labeled NMR samples (Fig. 1). Second, we carried NMR structure calculation. Our NMR structure depicts fold reminiscent of profilins, with eight strands interlinked by loops connecting three helices (Fig. 2). The relative positions and length of the helices and loops are comparable with Loki profilin-1, 2 and canonical eukaryotic profilins. The main difference to the Loki-1, 2 structures described recently was the absence of an extended loop called the Loki-loop^*13*^ and the presence of the extended N-terminal region. A detailed structural comparison with previously determined Lokiarchaeota- and a previously determined Heimdallarchaeota profilins reveal that Thorarchaeota (*TFG12995*.*1*) profilin (thorProfilin) is divergent to the Loki profilin-1^*13*^ (root mean squared deviation (RMSD) 3.29 Å) with the presence of the extended N-terminal region and a shorter loki-loop, and to the Heimdallarchaeota LC3 profilin (RMSD) 2.9 Å). Main, differences include the absence of an additional helix between residues 123-129, a parallel N and C terminal helix. However, the long N-terminal extension (residues 1-20) (Fig. 2) is also present in the Heimdallarchaeota LC3 profilin. Comparing to the human profilin-1, and apart from the N-terminal extension, major differences are seen between residues K5-Y59 and S91-A95. The polyproline binding motif corresponds to residues V24 (W3), Y28 (Y6), T29 (N9), W53 (W31), K139 (H133), L141 (L134), and I45 (Y139). Residues in brackets represent those in humanProfilin-1 (Fig. 3).

**Figure 1.**
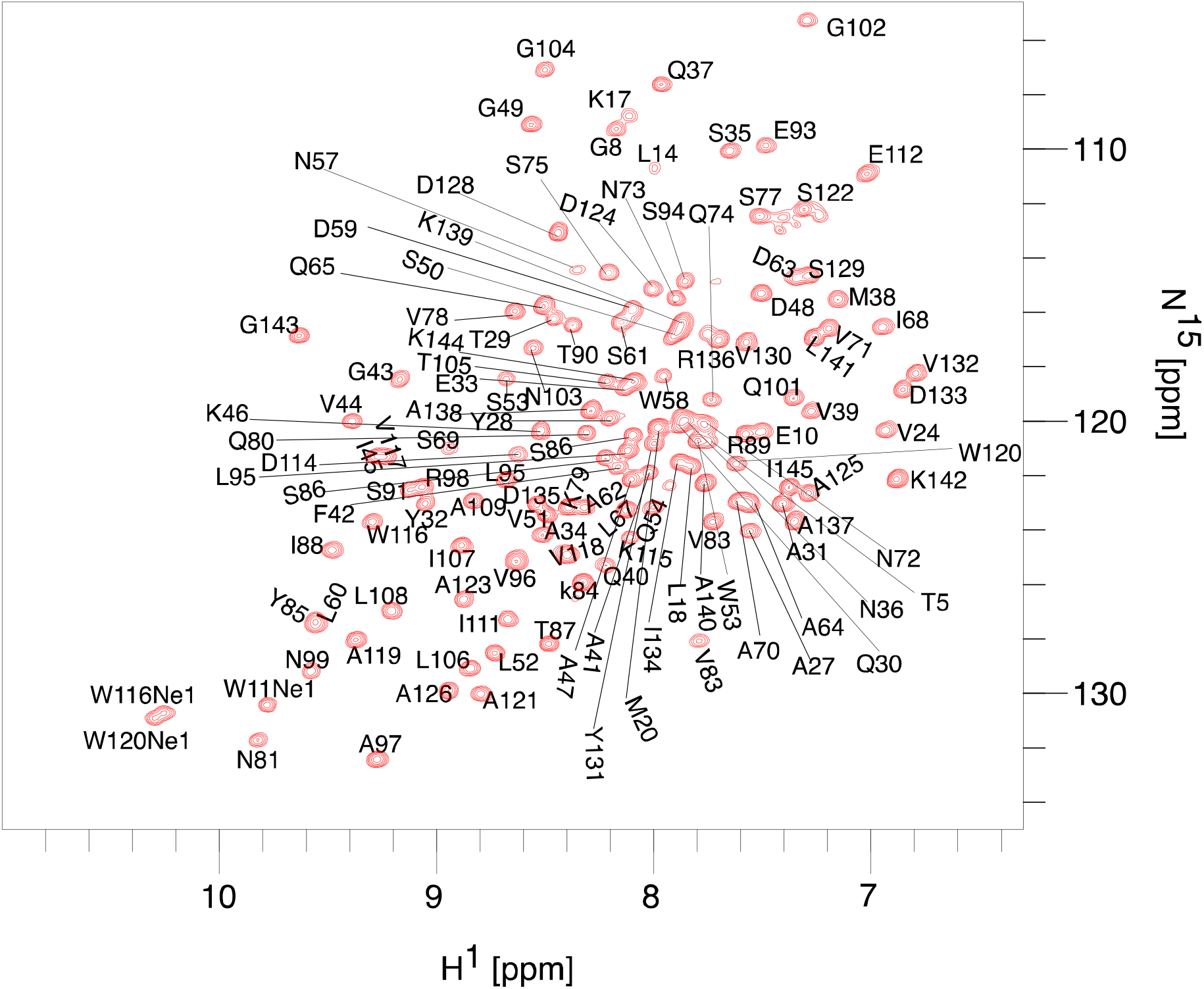
^1^H-^15^N TROSY correlation spectrum of Thorarchaeota profilin showing assigned atoms. A near complete backbone assignment of thorProfilin was carried out except for a few residues in the N-terminal region. Side-chains of Glutamine and Asparagine are not assigned or shown.

**Figure 2.**
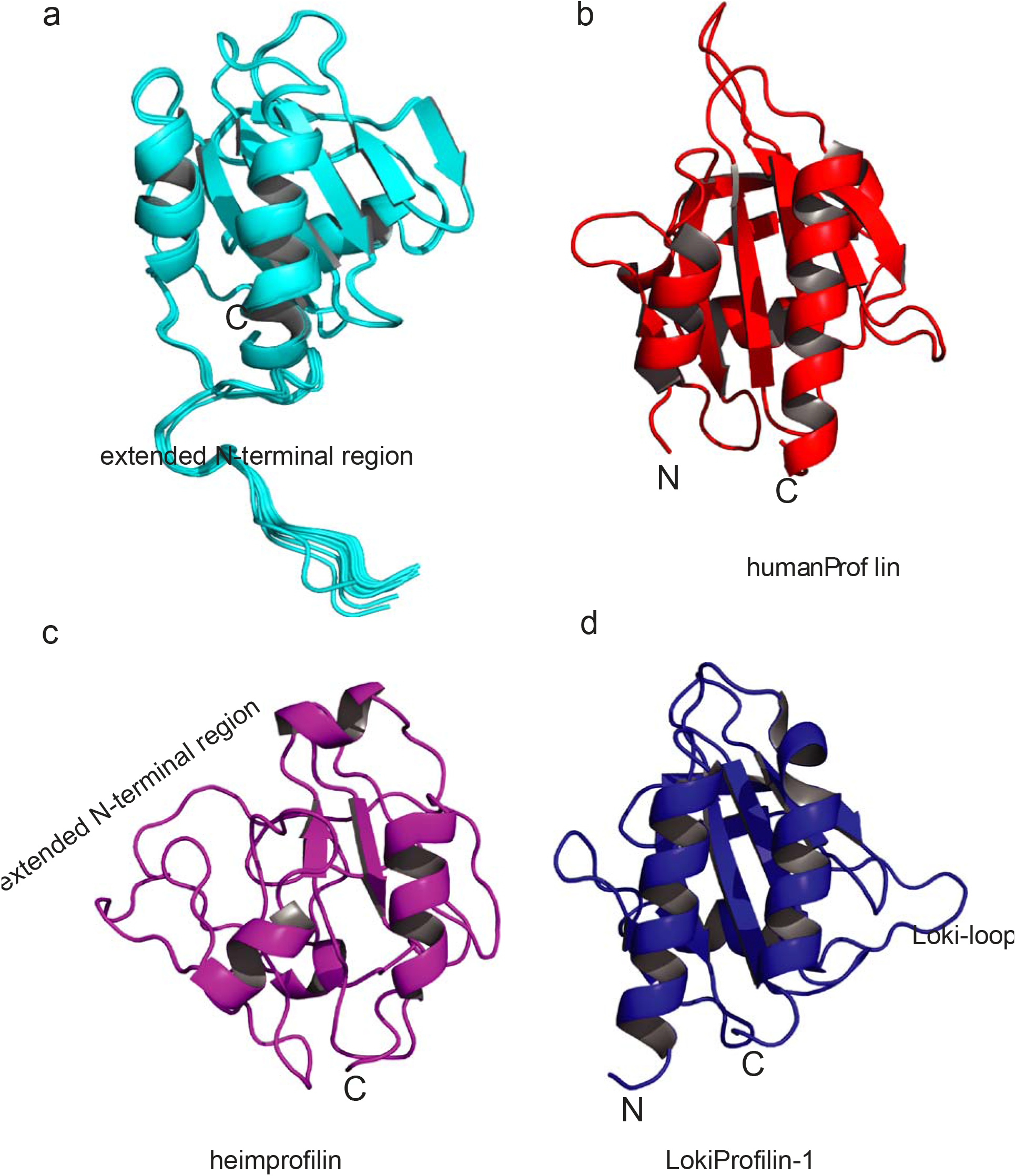
Thorarchaeota encodes profilin with an extended N-terminus. **a**, Structural representation of ThorProfilin. The extended N-terminus is displayed. **b**, Structural representation of human profilin-1 (1fil) reoriented in a similar fashion for comparison to the thorProfilin in (a) **c**, Structural representation of heimprofilin (PDB ID:6YRR) showing the extended N-terminal region. **d**, Structural representation of Loki profilin-1 (PDB ID:5zzb) showing the Loki-loop. The structural statistics are given in table 1. The structural coordinates have been deposited in the Protein data bank with PDB ID:7PBH

**Figure 3.**
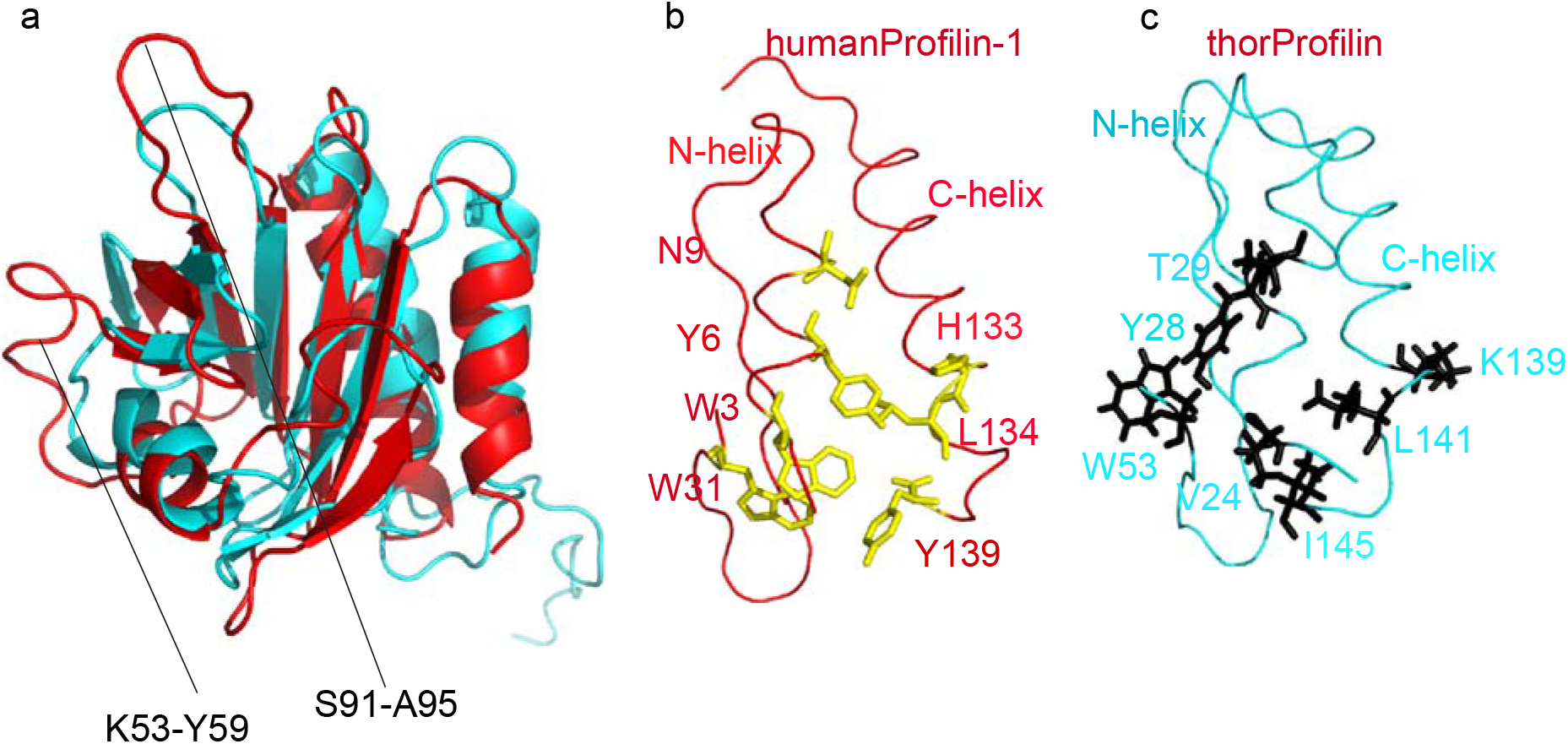
HumanProfilin-1 display similar polyproline binding site. **a**, overlay of humanProfilin-1(pdb:1fil) and thorPropfilin determined in this study. The major differences are indicated and corresponds to residues K5-Y59 and S91-A95. **b**, humanProfilin in a ribbon representation showing the polyproline binding residues. **c**, thorProfilin in a ribbon representation showing corresponding polyproline binding residues.

### Thorarchaeota profilin has a rigid core with a disordered N-terminus

In order to verify the dynamic nature of the N-terminal region, we determined the NMR relaxation parameters of the thorProfilin. We measured and evaluated the NMR R1, R1ρ, R2 and heteronuclear NOE (hetNOE) for the backbone amides ([^1^H]-^15^N). These NMR parameters report on motions in the ps-ms dynamics. Elevated values in R2 not seen in R1ρ could reflect exchange contributions in us-ms motions (faster than ca. 5 ms) while relative low hetNOE values below 0.6 will generally depict motions in ps-ns range. The NMR dynamic data determined for thorProfilin indicates that the residues between 22-145 are relative rigid and likely experiencing no slow motions less that ca. 5 ms. This observation is based on the fact that R2 and R1ρ contain almost identical contributions. On the other hand, residues 1-21 contain hetNOE values lower than 0.6 and a concomitant lower R2 values, indicative of motions in ps-ns (Fig. 4). These observation is in agreement with what was seen for heimProfilin, which contained a relative rigid protein central core and a disordered N-terminal (residues 1-24) ^*14,15*^.

**Table 1.**
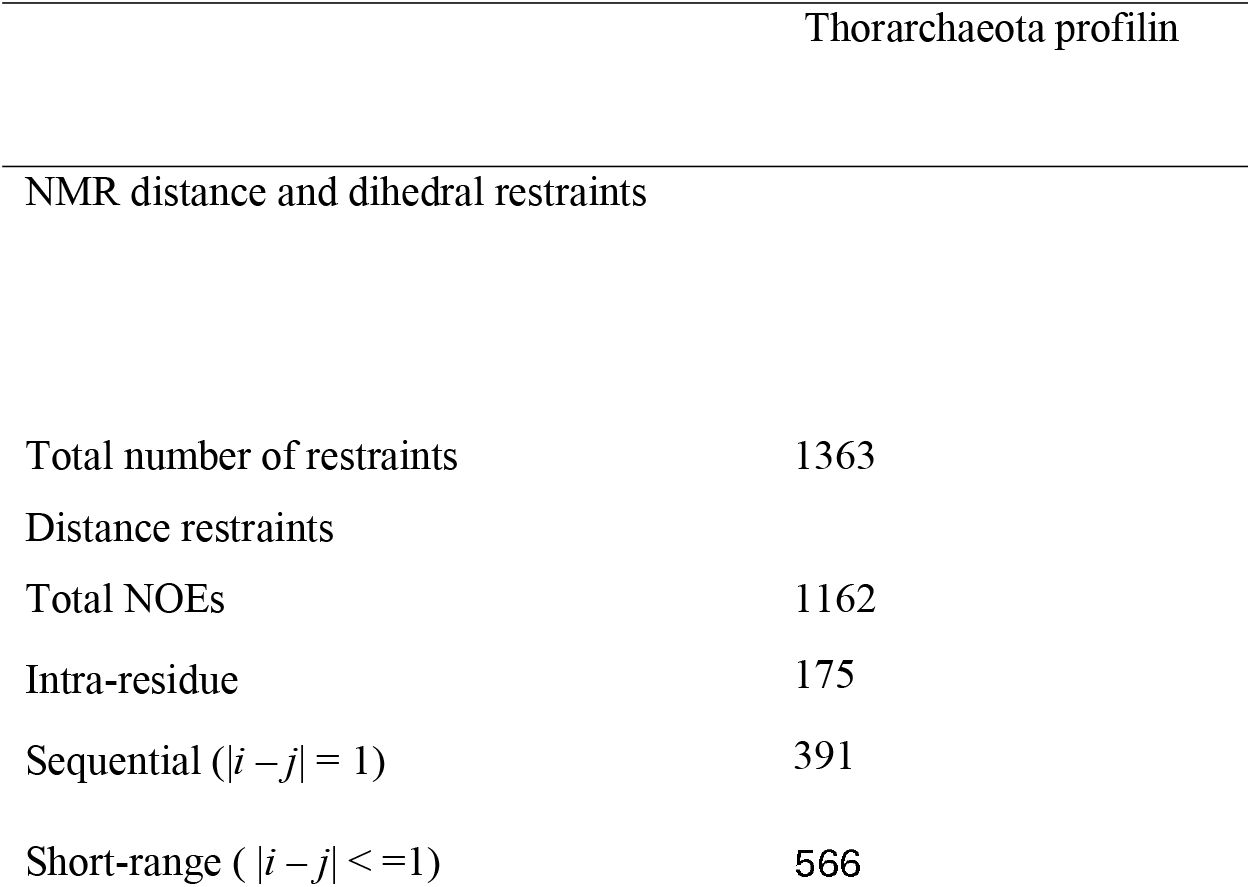
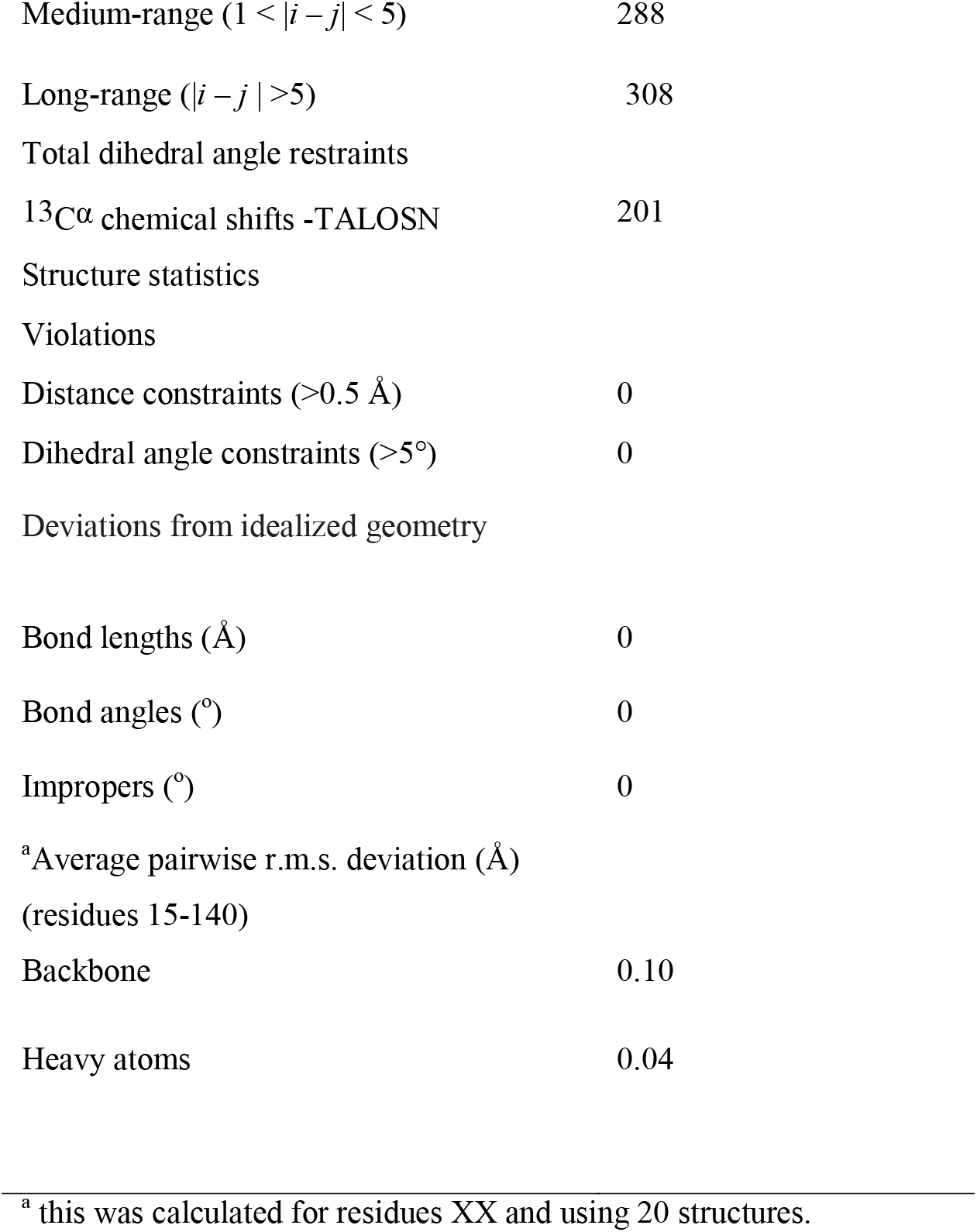
Nuclear magnetic resonance spectroscopy structural statistics

**Figure 4.**
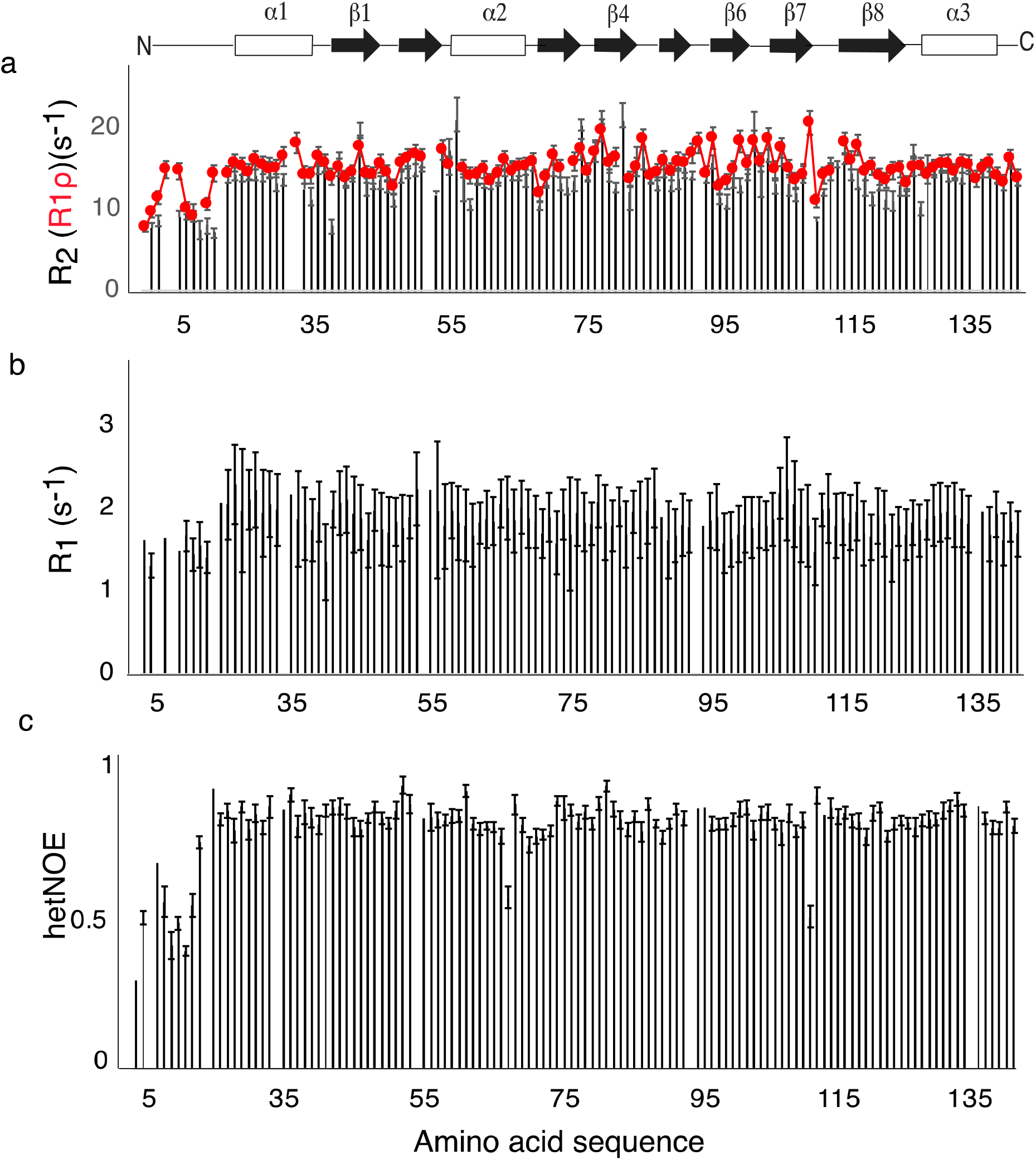
NMR backbone ([^1^H-]^15^N) dynamic characterization of. **a**, Transverse ^15^N R2 (R1ρ) relaxation rates versus the amino acid sequence **b**, Longitudinal ^15^N R1 relaxation rates plotted as a function of amino acid sequence. **c**, [^1^H]^15^N-heteronuclear NOE data (hetNOE) plotted as function of the amino acid sequence. The identical nature between the rates obtained for R2 and R1ρ indicates the absence of motions slower that ca 5ms. hetNOE on the other hand indicates ps-ns motions for the N-terminal residues. Overall, the backbone relaxation parameters indicate a very rigid molecule between residues 22-145. A cartoon representation of the positions of the secondary structural elements are indicated above (a). errors indicated by error bars are determined from the curve fit.

### Florescence microscopy reveals that thorarchaeota profilin co-localizes with eukaryotic actin cytoskeleton in cultured HeLa cells

Profilins from Loki-1 and 2, heimdall have been shown to regulate actin polymerization in *in vitro* pyrene-actin polymerization assay^*13*^. In addition, it was previously shown that thorProfilin binds to polyproline from Enabled/vasodilator stimulated phosphoprotein (Ena/VASP) family of proteins^*14*^. To verify if the thorarchaeota profilin might have any functional role *in vivo*, we monitored the localization eGFP-tagged thorProfilin with florescence microscopy and compare that to F-actin spatial location. To do this we used HeLa cells transfected with a plasmid expressing eGFP-thorprofilin. 24 hours post transfection, the HeLa cells were fixed and endogenous actin was stained with florescent CellMask™ F-actin marker. Thorproflin-F-actin localization was then monitored by observing the florescence of eGFP and F-actin in a confocal microscope with excitation at 488nm and emission at 565nm. We observed that GFP-thorProfilin co-localizes with endogenous F-actin in cultured HeLa cells (Fig. 5). These results indicate that the thorarchaeota profilin from *TFG12995*.*1* co-localizes with eukaryotic actin in a eukaryotic cell in addition to previously polyproline binding. Taken together, these results show that the thorProfilin and eukaryotic profilins likely have a common origin and highlights the complex origin of the eukaryotic cell.

**Figure 5.**
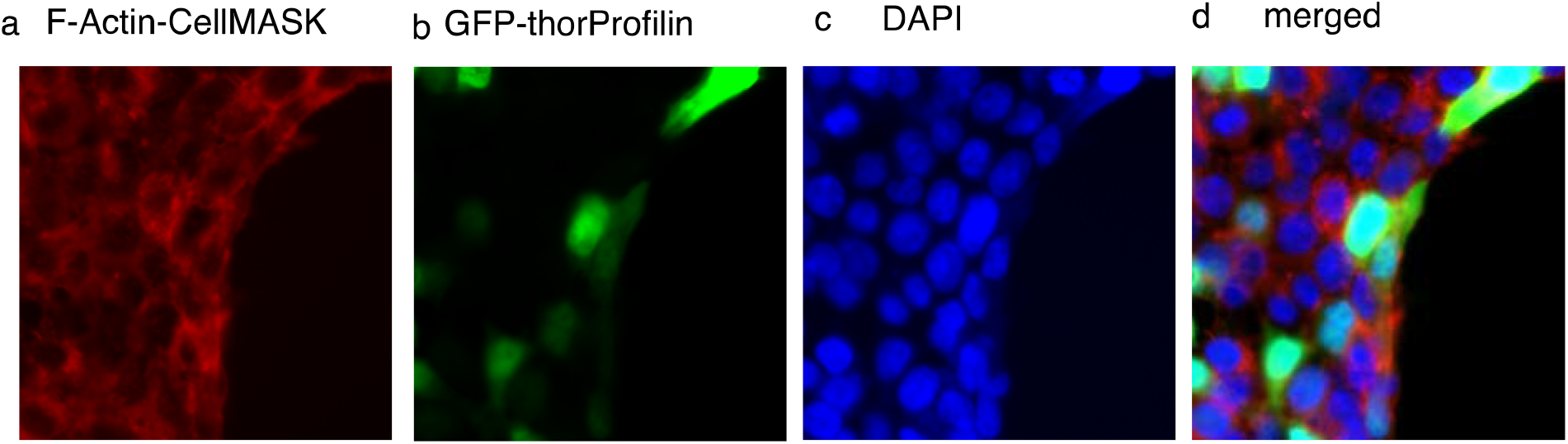
Fluorescence and co-localization of thorProfilin and F-actin in Hela cells. **a**, F-actin-cellMask indicating the location filamentous actin (red). **b**, eGFP-thorProfilin indicating over expression of thorProfilin (green). **c**, DAPI stain showing the nucleus (blue). **d**, a merger of a), b) and c) indicating the co-localization of F-actin and thorProfilin. The more yellow color indicates region where both F-actin and thorProfilin are co-localizing.

## DISCUSSION

It is now well-known that the Asgard superphylum encodes eukaryotic signature proteins that regulate many processes important for membrane maintenance and function^*5-7,13*^. The actin superfamily and associated proteins such as profilin, gelsolin and vasodilator-stimulated protein (VASP) are one such class of proteins that regulate cell cytoskeleton and are important in maintaining shape and motility of the cell. Actin is a cytoskeletal protein whose polymerization drives membrane remodeling via filament nucleation and elongation. Precisely, profilins from Lokiarchaeota, Odinarchaeota, and Heimdallarchaeota have been shown to regulate actin polymerization in pyrene-actin polymerization assay *in vitro*. In addition, it has been shown that VASP can regulate this actin polymerization reaction by interacting with profilins only from certain members of the Heimdallarchaeota and Thorarchaeota^*14*^. Moreover, profilins from all members of the Asgardarchaeota seem to interact with phospholipids^*13,14,16*^: a component of the eukaryotic membrane, indicating these Asgards might possess membrane organization similar to eukaryotes. Recently gelsolins from the Thoracrhaeota have been shown to regulate eukaryotic actin polymerization as well as co-localized with actin in cultured eukaryotic cells^*17*^. This means that Thorachaeota likely possesses eukaryotic actin regulatory characteristics. Here, we have determined the three-dimensional structure of Thorarchaeota *TFG12995*.*1* profilin and show that it contains an extended N-terminal extension similar to that seen in Heimdallarchaeota LC3. NMR backbone dynamic parameters determined for thorProfilin, revealed a protein with rigid central core and flexible N-terminus. In addition, we determined that thorProfilin co-localizes with Factin in cultured HeLa cells. Our findings thus indicate that some Asgardean profilins possesses characteristics and function analogues to eukaryotic profilins; actin polymerization regulation, polyproline binding, phospholipids binding, showing that the Asgardean cell already contained a great degree of complexity as compared to the present-day eukaryotes.

## Methods

### Protein expression and purification

The Thorarchaeota profilin used in the study was similar to that previously described^*14*^. Briefly, the gene was ordered from geneScript in pET28a vector with 6xHistidine-lipolyl tag and a TEV protease cleavage site at the N-terminus. Recombinant proteins were overexpressed in *E. coli* BL21*. At the start, the cells were grown at 37 °C in LB broth media. Protein expression was induced with 1 mM IPTG when the optical density (OD _600_) was 0.6 - 0.8. After induction, the cells were grown overnight at 25 °C. The cells were harvested by centrifugation and the cell pellet was re-suspended in purification buffer containing 25 mM Tris-HCl pH 8.0, 0.20 M NaCl, 1 mM DTT, 10 mM imidazole, 0.1% triton x100.

### Protein Expression for NMR

Thorprofilin for NMR experiments was expressed and purified as follows. After an initial growth of the cells in LB up to an OD_600_ of 0.6, the cells were then harvested by centrifugation at 4,000 x g for 5 min, rinsed and re-suspended in M9 medium (1g/L ^15^N-ammonium chloride and 1g/L ^13^C glucose). The cells were then allowed to grow for an additional 1 hour. Protein expression was induced by the addition of 1 mM IPTG overnight at 25 °C. For selective amino acid depletion, the M9 media was prepared as above, however, 1 g/L of the following amino acids (alanine, serine, isoleucine or leucine) were added respectively, 1 hour prior to addition of IPTG. Cells were thereafter harvested by centrifugation and the cell pellet was re-suspended in the binding and purification buffer (25 mM Tris-HCl pH 7.5, 0.20 M NaCl, 10 mM imidazole, 1 mM DTT, 0.1% Triton X100). Cells were lysed by sonication and the cell debris were separated from the soluble proteins by centrifugation at 45,000 x g for 60 min. The supernatant was filtered through a 0.45 μm and then 0.2 μm filter and thereafter loaded onto Nickel charged Sepharose column pre-equilibrated with purification buffer. After wash, the bound proteins were eluted with a buffer containing 400 mM Imidazole. The eluted proteins were desalted on a PD10 column (GE Healthcare). The lipoyl-tagged was cleaved by incubating with TEV protease overnight at RTP. The tag was removed by reloading the protein solution onto the Nickel charged column equilibrated with the binding buffer without Triton X100. The pure proteins were concentrated using a 3000-high molecular weight cutoff centrifugal filter (Merck-Millipore). The concentrated proteins were then subjected to size-exclusion chromatography using a Superdex-75 GL column (GE healthcare pre-equilibrated with 25 mM Tris-HCl 6.8, 150 mM NaCl. Purified proteins were pooled, concentrated, and stored at -20 °C until further use. Protein identity were checked on an SDS-PAGE and confirmed by MALDI mass spectrometry.

### Nuclear magnetic resonance spectroscopy experiments and assignment strategy

NMR experiments were carried out on Bruker spectrometer equipped with tripled resonance cryogenic probes operating at a proton larmor frequency of 600 MHz. The following 2D (^1^H-^15^N TROSY, ^1^H-^13^C HSQC) and 3D (TROSY-HNCACB, TROSY-HNCA, TROSY-HN(CO)CACB, HBHA(CO)HN, HCC(CO)HN) experiments were used for backbone and side-chain assignment. Assignments were confirmed by running ^1^H-^15^N TROSY and 2D HNCO experiments on samples made with specific amino acid depletion (see protein expression for NMR). All protein samples were either single labeled ^15^N, or double labeled ^15^N, ^13^C, at concentrations 3 mM in 25 mM sodium phosphate pH 6.8 supplemented with 3% D_2_O and 0.03% sodium azide. R2, R1, R1ρ and heteronuclear NOE were measured for thorProfilin in an interleaved manner with pulsegram from bruker library. For R2, R1, R1ρ, the relaxation delays were sampled for 8-10 delayed-duration in a pseudo-randomized manner. The relaxation delay was set to 1.2 s for R2, 1.5 s for R1 and 3 s for R1ρ. For the [^1^H]-^15^N-hetNOE experiment and reference experiment, the ^1^H saturation time was set to 3 s. All experiments were processed with Bruker TopSpin software and analyzed with the ccpNmr analysis program^*18*^ and bruker DynamicCenter2.5.3.

### NMR Structure Determination

NMR distance restraints were determined from 3D ^1^H-^1^H NOESY resolved in ^13^C-^1^H and ^15^N-^1^H TROSY experiments measured with the following specifications: 80 ms mixing time and 128 (^15^N or ^13^C) × 256 (^1^H) × 2048 (^1^H, direct). Structure calculations were done using the CYANA 3.98.13^*19*^ package as was described^*14*^. Briefly, the NOESY cross peaks were converted into upper distance restraints in an automated process in CYANA. These distance restraints in addition to the dihedral angles determined from backbone chemical shifts using TALOS-N^8^ and were then used as input for the structure calculations. The structures were calculated with 30,000 torsion angle dynamics steps for 100 conformers starting from random torsion angles by simulated annealing. For representation and analysis, the 20 conformers with the lowest target function values were selected. The structural statistics together with all input data for the structure calculations are presented in Table 1. The structural coordinates have been deposited in the protein data bank with PDB ID:7PBH

### Hela cell culture and transfection

HeLa cells (CCL-2) were cultured in DMEM media supplemented with 10% (v/v) heat-inactivated fetal bovine serum and 100 units/mL penicillin G and 100 μg/mL streptomycin solution (Gibco) in a humidified incubator at 37 °C with 5% CO2. 24 hours prior to transfection, the cells were seeded at 2 x 10 ^6 cells into a 24-well cluster plates with 1 mL of media per well. A glass coverslip was carefully placed at the bottom of each well prior to the seeding. For transfection, 1 μg of DNA was diluted into 100 μL of opti-MEM reduced serum media (Gibco) followed by addition of a μg PEI. The mixture was vortexed and incubated for 15 mins at room temperature. 100 μL of this DNA:PEI complex was then added dropwise to the well. The plate was gently agitated sidewise, and returned to the 37 °C incubator. Transfected cells were maintained for another 24 h.

### Florescence confocal microscopy

4 hours before imaging, cells were taken out of the incubator washed 3 times with cold PBS. The glass slides were then removed and placed on a parafilm for fixation. Cells were then fixed using paraformaldehyde thus: 4 % of 100 uL paraformaldehyde in PBS was added to the cells are allowed to incubate for 15 mins at RTP. PBS was then washed 3 times and 1 times CellMask^™^ orange Actin Tracking Stain (Thermofisher; Catalogue number: A57244) and allowed to incubate for another 15 mins at RTP to stain for F-actin. This was then washed 3 times with PBS and 1 times DAPI was added and incubated for 2 mins at RTP to stain for the nucleus. The glass slide was then mounted on a microscopic slide using a mounting solution. This was then allowed to stand for 1-2 hours at 4 °C. Cells were imaged using a LSM 710 Elyra S.1, AxioObserver confocal microscope with a Plan-Apochromat 63×/1.40 oil objective lens. Cells were imaged using the instrument’s, eGFP 488 nm, and 545 nm fluorophore default settings for F-actin stained with CellMask™ orange Actin Tracking Stain, respectively. images were processed with Imagej^*20*^.

## Funding Sources

The authors declare no competing financial interests have been declared. This work was supported by Wenner-Gren Stiftelserna fellow’s grants, Ake Wiberg, Magnus Bergvall and O.E Edla Johannsson foundation grants to C.C.

## Acknowledgements

This study made use of the NMR Uppsala infrastructure, which is funded by the Department of Chemistry - BMC and the Disciplinary Domain of Medicine and Pharmacy.

## Author Contributions

C. C designed research. R.I., M. D, S.L and C.C performed all research. C.C. wrote the paper. All authors have approved to the final version of the manuscript.

C. C. conceived study and performed all NMR. R.I performed cell culture, S.L and M.D performed microscopy. C. C. wrote the paper with contributions from all other authors. Funding: This work was supported by Wenner-Gren Stiftelsen Fellow’s Grants, Ake Wiberg, Magnus Bergvall and O.E Edla Johannsson foundation grants to C. C. This study made use of the NMR Uppsala infrastructure, which is funded by the Department of Chemistry - BMC and the Disciplinary Domain of Medicine and Pharmacy as well as the Imaging facility at Stockholm University. Conflicts of interest/Competing interests: The authors declare no conflict of interest. Ethics approval: Not applicable. Consent to participate: Not applicable. Consent for publication: All authors read and approved the manuscript. Availability of data and material: All data and material are available and can be obtain from the authors.

## Notes

### Competing Interest Statement

The authors have declared no competing interest.

